# Assessing GPT-4 for cell type annotation in single-cell RNA-seq analysis

**DOI:** 10.1101/2023.04.16.537094

**Authors:** Wenpin Hou, Zhicheng Ji

## Abstract

Cell type annotation is an essential step in single-cell RNA-seq analysis. However, it is a time-consuming process that often requires expertise in collecting canonical marker genes and manually annotating cell types. Automated cell type annotation methods typically require the acquisition of high-quality reference datasets and the development of additional pipelines. We assessed the performance of GPT-4, a highly potent large language model, for cell type annotation, and demonstrated that it can automatically and accurately annotate cell types by utilizing marker gene information generated from standard single-cell RNA-seq analysis pipelines. Evaluated across hundreds of tissue types and cell types, GPT-4 generates cell type annotations exhibiting strong concordance with manual annotations and has the potential to considerably reduce the effort and expertise needed in cell type annotation. We also developed GPTCelltype, an open-source R software package to facilitate cell type annotation by GPT-4.

## Main

In single-cell RNA-sequencing (scRNA-seq) analysis^1,2^, cell type annotation is a fundamental step to elucidate cell population heterogeneity and understand the diverse functions of different cell populations within complex tissues. Standard single-cell analysis software, such as Seurat^3^ and Scanpy^4^, routinely employ manual cell type annotation. These software tools assign single cells into clusters by cell clustering and conduct differential analysis to identify differentially expressed genes across cell clusters. Subsequently, a human expert compares canonical cell type markers with differential gene information to assign a cell type annotation to each cell cluster. This manual annotation approach requires prior knowledge of canonical cell type markers in the given tissues and is often laborious and time-consuming. Although several automated cell type annotation methods have been developed^5–13^, manual cell type annotation using marker gene information remains widely used in scRNA-seq analysis^14–28^.

Generative Pre-trained Transformers (GPT), including GPT-3, ChatGPT, and GPT-4, are large language models trained on massive amounts of data and capable of generating human-like text based on user-provided contexts. Recent studies have demonstrated the competitive performance of GPT models in answering biomedical questions^29–32^. Thus, we hypothesize that GPT-4, one of the most advanced GPT models, has the ability to accurately identify cell types using marker gene information. GPT-4 will potentially transform the manual cell type annotation process into a fully automated or semi-automated procedure, with optional help from human experts to fine-tune GPT-4-generated annotations (Figure 1a). Compared to other automated cell type annotation methods that require building additional pipelines and collecting high-quality reference datasets (Supplementary Table 1), GPT-4 offers cost-efficiency and seamless integration into existing single-cell analysis pipelines, such as Seurat^3^ and Scanpy^4^. The vast amount of training data enables GPT-4 to be applied across a wide variety of tissues and cell types, overcoming the limitations of other automated cell type annotation methods restricted to specific reference datasets. Additionally, the chatbot-like nature of GPT-4 allows users to easily adjust annotation granularity and provide feedback for iterative answer improvement (Figure 1a-b)^31^.

**Figure 1.**
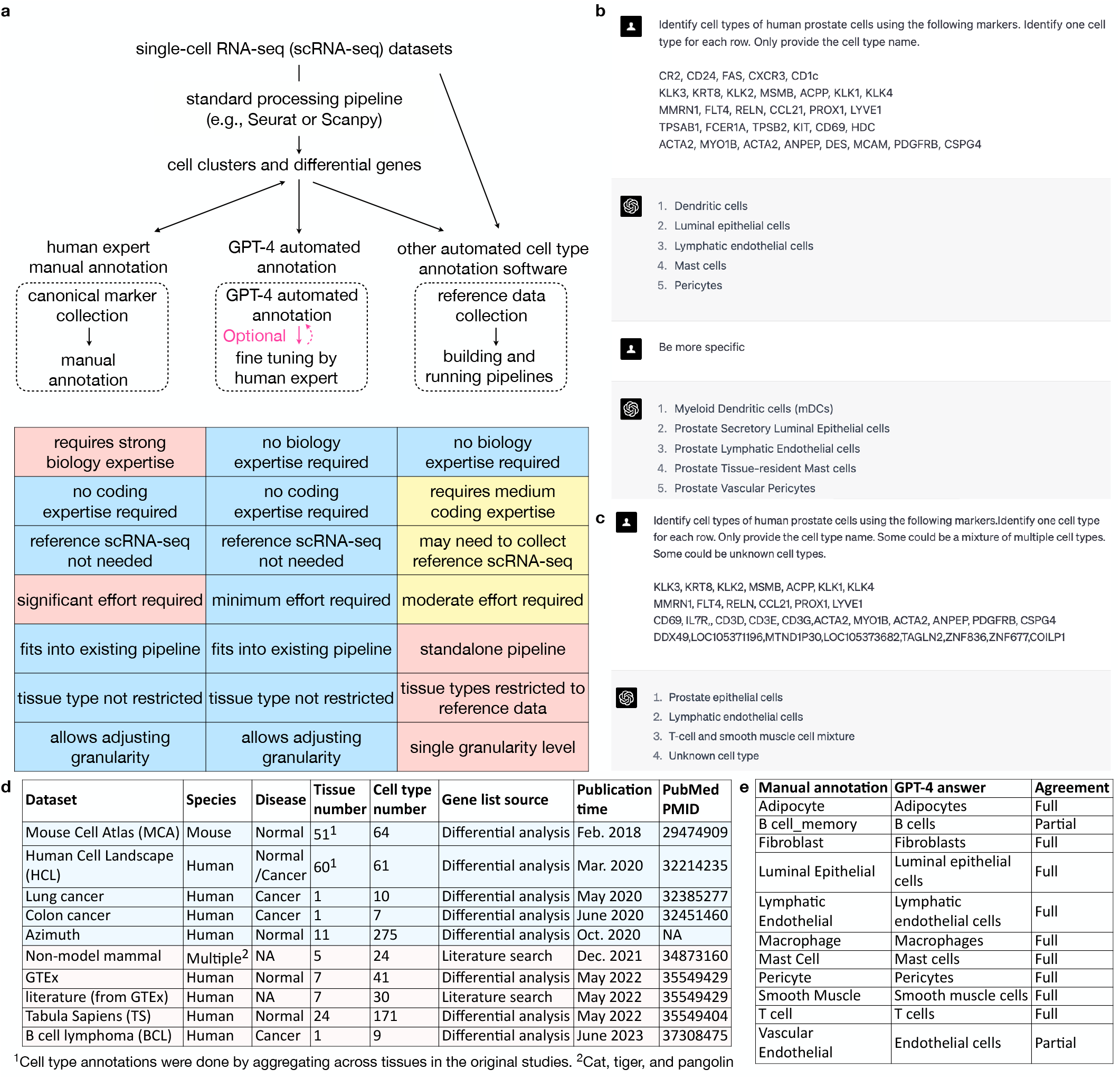
a, Diagram comparing cell type annotations by human experts, GPT-4, and other automated methods. **b**, An example showing GPT-4 prompts and answers for annotating human prostate cells with increasing granularity. **c**, An example showing GPT-4 prompts and answers for annotating single cell types (first two cell types), mixed cell types (third cell type), and new cell types (fourth cell type). **d**, Datasets included in this study. Datasets generated before Sep 2021, the cutoff date of GPT-4’s training corpus, were highlighted in blue, and others are highlighted in pink. **e**, Agreement between original and GPT-4 annotations in identifying cell types of human prostate cells.

In this study, we systematically assessed GPT-4’s cell type annotation performance across ten datasets^17–19,22,33–37^, hundreds of tissue types and cell types, in both normal and cancer samples, and in five species (Figure 1d). Computationally identified differential genes in eight scRNA-seq datasets and canonical marker genes identified through literature search in two datasets were used as inputs to GPT-4. Cell type annotation for HCL and MCA was performed and evaluated once by aggregating all tissues, similar to the original studies. In other studies, cell type annotation was performed and evaluated within each tissue. GPT-4 (June 13, 2023 version) was queried using GPTCelltype, a software tool we developed in this study (Methods). For competing methods, we evaluated GPT-3.5 (June 13, 2023 version), a prior version of GPT-4. We also evaluated other automatic cell type annotation methods of CellMarker2.0^38^, SingleR^39^, and ScType^40^ that provide references applicable to a large number of tissues (Methods, Supplementary Table 1). Cell type annotations by GPT-4 or competing methods were compared to manual annotations provided by the original studies. Specifically, each manually or automatically identified cell type annotation was assigned an unambiguous CL cell ontology name^41^ and a broad cell type name when applicable. A pair of manually and automatically identified cell type annotations was classified as “fully match” if they have the same annotation term or available CL cell ontology name, “partially match” if they have the same or subordinate (e.g., fibroblast and stromal cell) broad cell type name but different annotations and CL cell ontology names, and “mismatch” if they have different broad cell type names, annotations, and CL cell ontology names. Figure 1e shows an example of evaluating GPT-4 cell type annotations in a human prostate tissue literature search dataset. Details of all cell type annotations, their CL cell ontology names and broad cell type names, and agreements between manually and automatically identified cell type annotations are included in Supplementary Table 2.

The accuracy of cell type annotation may be influenced by the selection of input differential genes. We first assessed how the number of top differential genes obtained by the two-sided Wilcoxon test would affect the performance of GPT-4 cell type annotation (Supplementary Table 3). To facilitate comparison, we assigned agreement scores of 1, 0.5, and 0 to cases of “fully match”, “partially match”, and “mismatch” respectively, and calculated average scores within each dataset across cell types and tissues. Figure 2a shows that GPT-4 has the best agreement with human annotation when using the top 10 differential genes, and using more differential genes may reduce agreement. A plausible explanation is that human experts may only rely on a small number of top differential genes if they already provide a clear cell type annotation. We then evaluated the impact of different statistical methods for differential analysis on the performance of cell type annotation (Supplementary Table 3). The annotation performance marginally improves when differential genes are derived using the two-sided Wilcoxon test, as opposed to those obtained through the two-sided two-sample *t*-test (Figure 2a). This improvement could be attributed to the two-sided Wilcoxon test’s robustness, leading to differential genes more aligned with the specific cell type. In subsequent analyses, marker genes provided by Azimuth and literature search datasets as well as top 10 differential genes obtained from two-sided Wilcoxon test in all other datasets were used as inputs for GPT-4, GPT-3.5, and CellMarker2.0. SingleR and ScType were not performed on Azimuth and literature search datasets since full gene expression matrices were not available. Log-normalized gene expression matrices were used as inputs for SingleR and ScType in all other studies.

**Figure 2.**
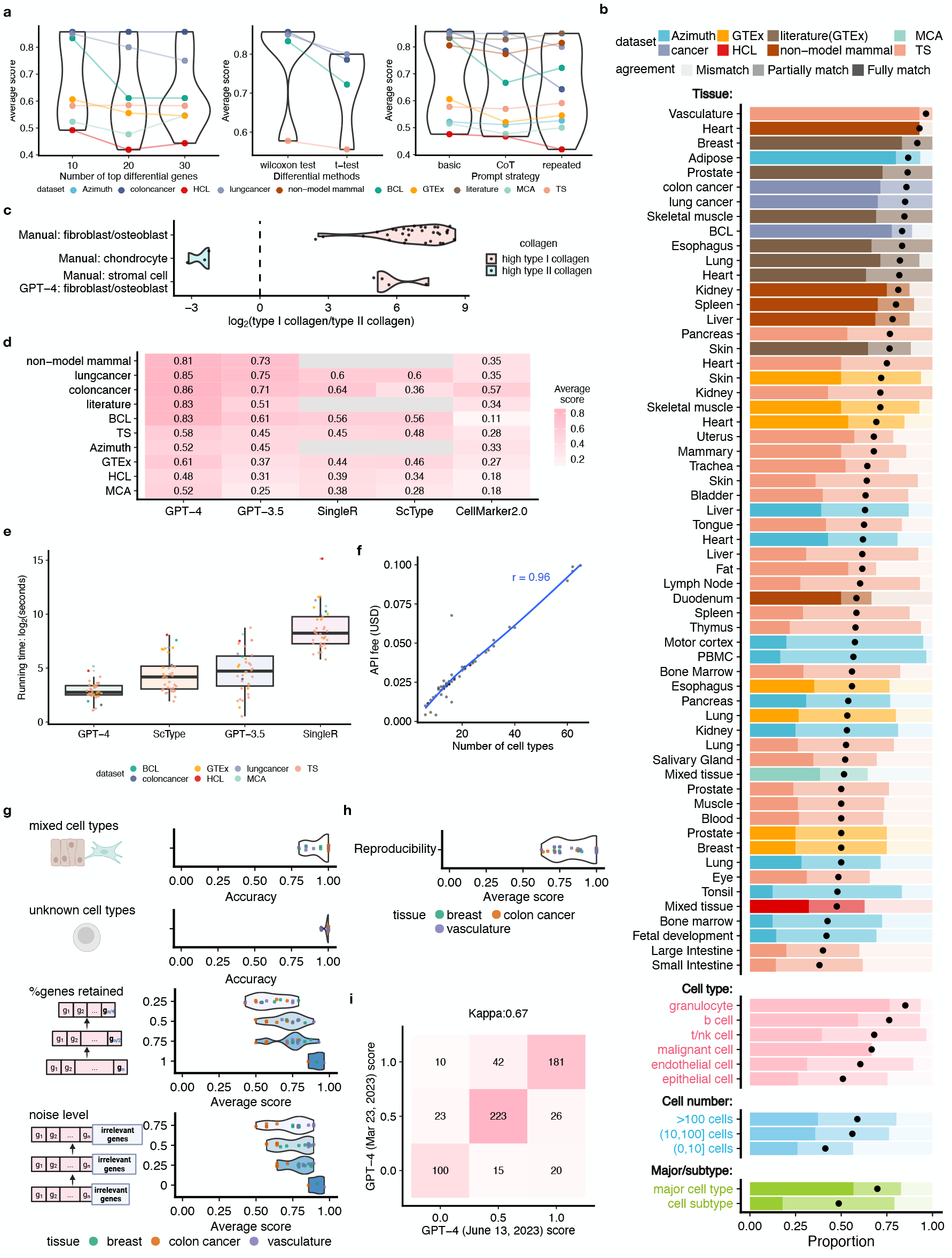
Performance evaluation of GPT-4 in cell type annotation. **a**, Average agreement scores with different numbers of top differential genes (left), with different statistical methods for differential analysis (middle), and with different prompt strategies (right). **b**, Proportion of cell types with different levels of agreement in each study and tissue, in top five most abundant broad cell types and malignant cells, in cell populations with different numbers of cells, and in major cell types or cell subtypes (from top to bottom). Average agreement scores are shown as black dots. **c**, log_2_-transformed ratio of averaged type I collagen gene expression (*COL*1*A*1,*COL*1*A*2) and type II collagen gene expression (*COL*2*A*1). **d-e**, Average agreement score (**d**) and running time (**e**) comparing different methods in each dataset. Each boxplot shows the distribution (center: median; bounds of box: 1st and 3rd quartiles; bounds of whiskers: data points within 1.5 IQR from the box; minima; maxima) of running time. **f**, The financial cost of querying GPT-4 API versus the number of cell types in each tissue and study. *r* represents Pearson correlation. **g**, Performance of GPT-4 identifying mixed and single cell types, known and unknown cell types, and with different levels of subsampling and noise. Each dot represents one round of simulation. **h**, Reproducibility of GPT-4 annotations. Each dot represents one cell type. **i**, Consistency of agreement scores between GPT-4 of June 13, 2023 version and GPT-4 of March 23, 2023 version. The numbers and colors in the plot represent the quantity of cell types from all relevant studies categorized accordingly.

The performance of GPT-4 can also be affected by how the input prompt message was structured. We tested a basic prompt strategy that only includes the necessary information, a prompt strategy inspired by chain-of-thought (CoT)^42^ that includes intermediate reasoning steps of an example, and a repeated prompt strategy where GPT-4 was queried multiple times and the most frequently appearing term was selected (Methods, Supplementary Table 3). Results show that the performances of different prompt strategies are comparable (Figure 2a). A potential reason is that cell type annotation does not involve complicated reasoning steps and the results from GPT-4 are highly reproducible. For simplicity, we used the basic prompt strategy for GPT-4 and GPT-3.5 in subsequent analysis.

In almost all studies and tissues, GPT-4 annotations fully or partially match manual annotations for at least 75% of cell types (Figure 2b), demonstrating GPT-4’s ability to generate cell type annotations comparable to those of human experts. Notably, the agreement is especially pronounced for marker genes identified by literature searches, where GPT-4’s annotations fully match with manual annotations in 70% of cell types in most tissues. While there is a decrease in agreement for marker genes identified by differential analysis, the agreement remains commendably high, rendering GPT-4 suitable for a broad range of datasets. It is worth noting that the Azimuth, HCL, MCA, lung cancer, and colon cancer datasets were published before September 2021 (Figure 1d), the cutoff date of GPT-4’s training corpus. Thus, the results of these datasets should be interpreted with care since they may not fully reflect the performance of GPT-4 when dealing with new gene sets not in the training corpus. Figure 2b also shows the performance of GPT-4 in the five most abundant broad cell types and malignant cells across all studies. GPT-4 shows higher agreement in immune cell types such as granulocytes compared to other cell types such as epithelial cells. GPT-4 was able to identify malignant cells in colon and lung cancer datasets, but failed in the BCL dataset. A potential reason is that there is not a well-defined signature gene set for B lymphoma cells, making the cell type hard to identify. The identification of malignant cells could benefit from other approaches such as copy number variation^43^, as in the original study^34^. In smaller cell populations comprising no more than 10 cells, GPT-4 exhibits a modest reduction in performance (Figure 2b), which might be attributed to the limited information available in these rare cell groups. Finally, we classified each manually identified cell type annotation as a major cell type (e.g., T cells) or a cell subtype (e.g., CD4 memory T cells). GPT-4 has a significantly higher proportion of “fully match” cases in major cell types, although there are still more than 75% cases of “fully match” and “partially match” combined for cell subtypes (Figure 2b).

The low agreement between GPT-4 and manual annotations in some cell types does not necessarily imply that GPT-4 annotation is incorrect. For instance, cell types classified as stromal cells include fibroblasts and osteoblasts, which express type I collagen genes, as well as chondrocytes, which express type II collagen genes. For cells manually annotated as stromal cells, GPT-4 assigns cell type annotations with higher granularity (e.g., fibroblasts and osteoblasts), resulting in partial matches and a lower agreement. For cell types manually annotated as stromal cells but annotated as fibroblasts or osteoblasts by GPT-4, type I collagen genes have significantly higher expression than type II collagen genes (Figure 2c). This agrees with prior knowledge and the pattern observed in cell types manually annotated as chondrocyte, fibroblast, and osteoblast (Figure 2c), suggesting that GPT-4 provides more accurate cell type annotations than manual annotations for stromal cells.

We next compared the performance of GPT-4 with other competing methods by averaging the agreement scores within each study across all tissues and cell types (Figure 2d). In all datasets, GPT-4 performs significantly better than other methods, including its prior version GPT-3.5. GPT-3.5 has similar performance compared to SingleR, ScType and CellMarker2.0. Using GPTCelltype as an interface and working directly on the identified differential genes, GPT-4 is also significantly faster than other methods (Figure 2e). GPT-4’s efficiency is partly attributed to its use of differential genes from standard single-cell analysis pipelines like Seurat^3^. Given the integral role of these pipelines in analyzing scRNA-seq datasets, we regard the differential genes as immediately available for GPT-4. In comparison, other existing methods such as SingleR and ScType need to take extra steps and reprocess the gene expression matrices, which can be computationally intensive. Note that we used an in-house implementation of ScType (Methods) with significantly increased computational efficiency since the original ScType implementation cannot process large datasets in reasonable time. We did not compare the running time of CellMarker2.0 since it requires users to manually input gene sets on its online user interface. Compared to other methods that are free of charge, GPT-4 requires a $20 monthly subscription fee for using the online web portal. The financial cost for using GPT-4 API is linearly correlated with the number of cell types in the query, and does not exceed $0.1 for all queries in this study (Figure 2f).

We further tested the performance of GPT-4 when dealing with more complicated situations in real data analyses (Figure 1c). We first tested GPT-4’s ability to identify a cell cluster representing a mixture of cell types, which may occur when a cluster contains a large number of doublets or has low-resolution cell clustering. We generated simulated datasets by combining canonical markers from two distinct cell types in half of the instances and using canonical markers from a single cell type in the other half (Methods). GPT-4 discriminated between single and mixed cell types with an average accuracy of 93% (Figure 2g). We then evaluated GPT-4’s ability to identify new cell types with marker genes not documented by existing literature. We created simulation datasets using randomly selected genes as cell type markers in half of the cases and canonical markers from a single cell type in the other half (Methods). GPT-4 is able to differentiate known and unknown cell types with an average accuracy of 99% (Figure 2g). We then tested the performance of GPT-4 with partial marker gene information by randomly subsampling 75%, 50%, and 25% of original marker genes (Methods). GPT-4’s performance decreases with a smaller number of marker genes but still maintains at a high level even with half of the original marker genes available (Figure 2g). Finally, we evaluated the performance of GPT-4 when the input information is contaminated by adding random genes to the marker gene list (Methods). GPT-4’s performance only decreases slightly with a higher level of contamination and maintains at a high level (Figure 2g). All these results suggest that GPT-4 has a robust performance when dealing with different situations.

We also evaluated the reproducibility of GPT-4 annotations leveraging results in previous simulation studies (Methods). On average, GPT-4 generated identical annotations for the same cell type markers in 85% of cases (Figure 2h), showing a high level of reproducibility. In addition, we compared the annotations between an older version of GPT-4 (March 23, 2023 version) and the June 13, 2023 version of GPT-4. In most instances, the two versions of GPT-4 exhibit identical agreement scores, achieving a Cohen’s Kappa of 0.65, which indicates a substantial level of consistency (Figure 2i).

In conclusion, we systematically assessed the performance of GPT-4 for cell type annotation in scRNA-seq datasets and found that there is a high level of agreement between cell type annotations generated by GPT-4 and by human experts. We also developed GPTCelltype that can be employed as a dependable tool for automated cell type annotation of single-cell RNA-seq data, substantially reducing the time and effort required for manual annotation. To enable a semi-automated process where human experts can interact with GPT-4 to further fine-tune the cell type annotations, GPTCelltype also provides the option to generate prompts that can be directly fed into GPT-4’s online user interface.

Although GPT-4 has a strong performance in cell type annotation and outperforms existing methods according to our assessment, users should still be aware of several limitations when applying GPT-4 for cell type annotation. First, unlike other cell type annotation methods, the training corpus of GPT-4 is largely undisclosed, making it difficult to explicitly verify the basis upon which GPT-4 generates annotations. Certain human effort may still be needed to critically evaluate the quality and reliability of the annotations generated by GPT-4. Second, the involvement of human experts in the optional fine-tuning step may negatively impact the repeatability of the results due to the added subjectivity and may reduce the scalability of the approach when applied to a large number of datasets. Third, a high level of noise in scRNA-seq data and unreliable differential genes may negatively impact GPT-4’s cell type annotations. Finally, predominantly relying on GPT-4 for cell type annotations could be risky in the case of AI hallucination. It is recommended that human experts confirm the validity of cell type annotations generated by GPT-4 before conducting downstream analyses.

## Methods

### Dataset collection

For the HuBMAP Azimuth project, manually annotated cell types and their marker genes were downloaded from the Azimuth website (https://azimuth.hubmapconsortium.org/). Azimuth provides cell type annotations for each tissue at different granularity levels. We selected the level of granularity with the fewest number of cell types, provided that there were more than 10 cell types within that level. Details of how marker genes were generated is not available.

For GTEx^17^ dataset, manually annotated cell types, differential gene lists, and gene expression matrices were downloaded directly from the publication^17^. In the original study, gene expression raw counts were library-size-normalized and log-transformed after adding a pseudocount of 1 with SCANPY^4^. ComBat^44^ was used to account for the protocol- and sex-specific effects with SCANPY^4^. Welch’s t-test was then performed to identify differential genes comparing one cell type against the rest. For each cell type, genes were ranked increasingly by p-values, and genes with the same p-values were further ranked decreasingly by t-statistics. Top 10, 20, and 30 differential genes were used in this study. Lists of marker genes through literature search and the corresponding cell types were downloaded from the same study^17^, and only cell types with at least 5 marker genes were used.

For HCL^19^ dataset, manually annotated cell types, differential gene lists, and gene expression matrix were downloaded directly from the publication^19^. In the original study, gene expression raw counts underwent a batch removal process to facilitate cross-tissue comparison, and were subsequently library size normalized and log-transformed after adding a pseudocount of 1. Two-sided two-sided Wilcoxon rank-sum test was then performed to identify differential genes comparing one cell type against the rest using Seurat^3^. Differential genes were further selected by log foldchange larger than 0.25, Bonferroni-adjusted p-value smaller than 0.1, and expressed in at least 15% of cells in either population. For each cell type, genes were ranked increasingly by p-values, and genes with the same p-values were further ranked decreasingly by two-sided Wilcoxon test statistics. Top 10, 20, and 30 differential genes were used in this study.

For MCA^18^ dataset, manually annotated cell types, differential gene lists, and gene expression matrix were downloaded directly from the publication^19^. In the original study, gene expression raw counts underwent a batch removal process to facilitate cross-tissue comparison, and Seurat^3^ was used to perform preprocessing and differential analysis. For each cell type, genes were ranked increasingly by p-values, and genes with the same p-values were further ranked decreasingly by log fold changes. Top 10, 20, and 30 differential genes were used in this study.

For non-model mammal dataset^37^, manually annotated cell types and lists of marker genes through literature search were downloaded directly from the original study.

For TS^33^, BCL^34^, lung cancer^36^, and colon cancer^35^ datasets, manually annotated cell types and raw gene expression count matrices were downloaded directly from original studies. Raw counts were normalized by library size and log-transformed after adding a pseudocount of 1. Seurat FindAllMarkers() function with default settings was used to obtain differential genes by comparing one cell type with the rest within each tissue. Briefly, genes with at least 0.25 log fold change between two cell populations and detected in at least 10% of cells in either cell population were retained. Two-sided Wilcoxon rank-sum test was then performed for differential analysis. In addition, two-sided two-sample *t*-test was also performed for differential analysis using the FindAllMarkers() function with default settings. For each cell type, genes were ranked increasingly by p-values, and genes with the same p-values were further ranked decreasingly by log fold changes. Top 10, 20, and 30 differential genes were used in this study.

### Cell type annotation methods

#### GPT-4 and GPT-3.5

All GPT-4 (June 13, 2023 version) and GPT-3.5 (June 13, 2023 version) cell type annotations in this study were performed using GPTCelltype, an R software package we developed as an interface for GPT models. GPTCelltype takes marker genes or top differential genes as input, and automatically generates prompt message using the following template with the basic prompt strategy:

“Identify cell types of TissueName cells using the following markers separately for each row. Only provide the cell type name. Do not show numbers before the name. Some can be a mixture of multiple cell types.*\*n GeneList”.

Here “TissueName” is a variable that will be replaced with the actual name of the tissue (e.g., human prostate), and “GeneList” is a list of marker genes or top differential genes. Genes for the same cell population are joined by comma (,), and gene lists for different cell populations are separated by the newline character (*\*n). GPT-4 or GPT-3.5 was then queried using the generated prompt message through OpenAI API, and the returned information was parsed and converted to cell type annotations.

For chain-of-thought (CoT) prompt strategy, the following sentence was added to the beginning of the message generated by the basic prompt strategy: “Because CD3 gene is a marker gene of T cells, if CD3 gene is included in the marker gene list of an unknown cell type, the cell type is likely to be T cells, a subtype of T cells, or a mixed cell type containing T cells.”

For repeated prompt strategy, GPT-4 was queried with the basic prompt strategy repeatedly for five times. The annotation result that appears most frequently among the five queries was selected as the final cell type annotation.

GPT-4 (March 23, 2023 version) cell type annotations were performed by manually copying and pasting prompt messages to GPT-4 online web interface (https://chat.openai.com/). The prompt message was constructed using the following template:

“Identify cell types of TissueName cells using the following markers. Identify one cell type for each row. Only provide the cell type name. *\*n GeneList”.

#### SingleR

SingleR^39^ (version 1.4.1) R packages was used to perform cell type annotations with default settings. For HCL and MCA datasets, the gene expression matrices after batch effect removal, library size normalization, and log-transformation across all tissues were used as input. For all other datasets, SingleR was performed separately within each tissue, and the input is the log-transformed and library size normalized gene expression matrix. The built-in Human Primary Cell Atlas reference^45^ was used as the reference dataset for all SingleR annotations. SingleR generates single-cell level cell type annotations by returning an assignment score matrix for each single cell and each cell type label in the reference. To convert single-cell level annotations to cell-cluster level annotations, for each manually annotated cell type, we assigned the reference label with assignment scores summed across all single cells in that manually annotated cell type as the predicted cell type annotation.

#### ScType

ScType^40^ (version 1.0) R packages was used to perform cell type annotations with default settings. The original implementation of ScType is computationally inefficient and cannot deal with large datasets in reasonable time. Thus, we implemented an in-house version of ScType that uses vectorization to optimize the most time-consuming steps, while still generating the exact same output of the original ScType software.

The input gene expression matrices to ScType were exactly the same as used in SingleR described above. The built-in cell type marker database was used as the reference for all ScType annotations. Manually annotated cell types were treated as cell clusters and given as inputs to ScType. ScType directly generates cluster-level cell type annotations.

#### CellMarker2.0

CellMarker2.0^38^ only provides an online user interface and does not have a software implementation. We used the exact same marker gene sets or top 10 differential gene sets used for GPT-4 and GPT-3.5 cell type annotations as inputs of CellMarker2.0.

### Simulation studies and reproducibility

To generate simulation datasets, we used canonical cell type markers through GTEx literature search of human breast cells, top 10 differential genes from the human colon cancer dataset, and top 10 differential genes from the vasculature tissue of the TS dataset as templates. Simulation studies were performed separately for each of the three tissue types.

To generate simulation datasets of mixed cell types, marker genes for each mixed cell type were created by combining the marker gene lists of two randomly selected cell types. Ten mixed cell types were generated in each simulation iteration. Additionally, we incorporated the original cell type markers of ten randomly chosen cell types as negative controls of single cell types. This entire simulation process was repeated five times. Subsequently, GPT-4 was queried using these simulated marker gene lists, and its performance in differentiating between mixed and single cell types was assessed.

To generate simulation datasets of unknown cell types, we compiled a list of all human genes using the Bioconductor org.Hs.eg.db package^46^. In each simulation iteration, ten simulated unknown cell types were generated. The marker genes for each unknown cell type were produced by combining ten randomly selected human genes. Additionally, we included ten real cell types and their marker genes as negative controls of known cell types, similar to the previous simulation study. This entire simulation process was repeated five times. Subsequently, GPT-4 was queried using these simulated marker gene lists, and its performance in distinguishing between known and unknown cell types was assessed.

To generate simulation datasets with partial marker gene information, we randomly subsampled 25%, 50%, or 75% of the original marker genes. The simulation process was repeated five times. Subsequently, GPT-4 was queried using these subsampled marker gene lists, and the performance was assessed by agreement scores.

To generate simulation datasets with contaminated information, we added randomly selected human genes to the original marker genes. The numbers of randomly selected genes are 25%, 50%, or 75% of the number of original marker genes. The simulation process was repeated five times. Subsequently, GPT-4 was queried using these subsampled marker gene lists, and the performance was assessed by agreement scores.

We assessed the reproducibility of GPT-4 responses by leveraging the repeated querying of GPT-4 with identical marker gene lists of the same negative control cell types in both simulation studies. For each cell type, reproducibility is defined as the proportion of instances in which GPT-4 generates the most prevalent cell type annotation. For instance, in the case of vascular endothelial cells, GPT-4 produces “endothelial cells” 8 times and “blood vascular endothelial cells” once. Consequently, the most prevalent cell type annotation is “endothelial cells,” and the reproducibility is calculated as 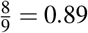.

### GPT-4 API financial cost

According to information provided by OpenAI, the API cost for running GPT-4 June 13, 2023 version is $0.03 for every thousand input tokens and $0.06 for every thousand output tokens. For each query, we obtained *i* and *o*, which represent the numbers of input tokens and output tokens respectively, through the OpenAI API. The total API financial cost is thus calculated as $(0.00003*i* + 0.00006*o*).

## Acknowledgments

Z.J. was supported by the National Institutes of Health under Award Number 1U54AG075936-01. W.H. was supported by the General Fund at Columbia University Department of Biostatistics and by the National Institute Of General Medical Sciences of the National Institutes of Health under Award Number R35GM150887. The content is solely the responsibility of the authors and does not necessarily represent the official views of the National Institutes of Health. The manuscript was polished by GPT-4.

## Author Contributions

All authors conceived the study, conducted the analysis, and wrote the manuscript.

## Data Availability

The data used in this manuscript are all downloaded from publicly available data sources. Specifically, HubMAP Azimuth data were downloaded from the Azimuth website (https://azimuth.hubmapconsortium.org/). GTEx manually annotated cell types and differential gene lists were downloaded from the supplementary materials of the original study^17^. GTEx gene expression matrix was downloaded from the GTEx website (https://gtexportal.org/home/datasets). Marker genes from literature search were downloaded from the supplementary materials of the original study^17^. HCL manually annotated cell types and differential gene lists were downloaded from the supplementary materials of the original study^19^. HCL gene expression matrix was downloaded from figshare (https://figshare.com/articles/dataset/HCL_DGE_Data/7235471). MCA manually annotated cell types and differential gene lists were downloaded from the supplementary materials of the original study^18^. MCA gene expression matrix was downloaded from figshare (https://figshare.com/s/865e694ad06d5857db4b). BCL gene expression matrix and manually annotated cell types were downloaded from Zenodo (https://zenodo.org/record/7813151). Colon cancer gene expression matrix and manually annotated cell types were downloaded from GEO under accession number GSE132465. Lung cancer gene expression matrix and manually annotated cell types were downloaded from GEO under accession number GSE131907. TS gene expression matrix and manually annotated cell types were downloaded from UCSC Cell Browser(https://cells.ucsc.edu/?ds=tabula-sapiens). Marker genes and cell type annotations for the non-model mammal dataset were downloaded from the supplementary materials of the original study^37^. All relevant information about data is described in the Methods section. All data generated in this study are included in the supplementary table.

## Code Availability

The GPTCelltype package (v.1.0.0) is provided as an open-source software package with a detailed user manual available in Github repository https://github.com/Winnie09/GPTCelltype. The software is released in Zenodo under the accession code DOI: 10.5281/zenodo.8317406 for all versions (https://doi.org/10.5281/zenodo.8317406)^47^. All codes to reproduce the presented analyses are publicly available in Github repository https://github.com/Winnie09/GPTCelltype_Paper and also in Zenodo under the accession code DOI: 10.5281/zenodo.8317410 (https://zenodo.org/record/8317410)^48^. R version 4.0.2 was used to perform the analyses in the manuscript.

